# Neural field patient-specific super resolution for enhanced 1.5 Tesla brain MRI visualization

**DOI:** 10.1101/2025.09.27.678985

**Authors:** Nicolas Gonzalez-Romo

## Abstract

Brain magnetic resonance imaging serves as a cornerstone of preoperative neurosurgical assessment. Neural fields represent an emerging machine learning approach capable of super-resolution reconstruction and novel view synthesis without requiring large training datasets. We evaluated ten 1.5-Tesla brain MRI sequences (nine anisotropic and one isotropic) to train patient-specific neural field models using a proprietary framework (Radscaler©). Image quality assessment was performed on reconstructions upscaled by factors of 2×, 3×, and 4× relative to original resolution. The method achieved favorable quality metrics across all scaling factors: mean SSIM of 0.85 (±0.04), MS-SSIM of 0.95 (±0.01), and LPIPS of 0.09 (±0.04). Neural field reconstruction enabled enhanced visualization of micro-anatomical structures through improved spatial resolution and interpolation of intermediate views not present in the original acquisition. These findings demonstrate that neural fields provide a clinically viable approach for volumetric MRI super-resolution and novel view synthesis, particularly valuable for addressing anisotropic acquisition limitations in neurosurgical planning.

## Introduction

Brain magnetic resonance imaging (MRI) is a routine study performed for the assessment of intracranial pathology. High magnetic field > 7 Tesla MRI (1–4) delivers ultra resolution images of the brain with increased signal to noise ratios, allowing detailed visualization of the gray white matter junction, mesial temporal lobe and brainstem anatomy. Powerful magnetic field superconductors however, are not available for everyday clinical practice in most settings, and the standard 1.5-Tesla brain MRI scan is still commonly performed for neurological diagnosis in developing countries. At the same time, low magnetic field <0.1-Tesla MRI scanners due to their lower cost and portability, promise to become a good alternative for resource limited and remote locations(5).

In order to improve quality of lower resolution MRI imagining, machine learning super resolution methods have been implemented. Compared to traditional bilinear and bicubic interpolation, machine learning algorithms such as convolutional neural networks(6), generative adversarial networks(7), and latent diffusion models(8) have demonstrated improvement of low resolution MRI image quality but these networks demand large datasets to learn a high resolution prior in order to up sample low resolution images at test time.

Neural fields, also know as implicit neural representations, are a more recent machine learning method hat given a set of images, can learn a 3D scene representation from input 2D images. The neural network learns a parametric field, where every coordinate of the Cartesian space is assigned a pixel color value in the weights of the model(9). This neural representation has four important properties. First, it can be queried to visualize images at arbitrarily increasing resolutions, without suffering from the traditional pixel artifacts present with other types of interpolation methods. Second, new images can be obtained from different viewpoints, not present in the training set, a task known as novel view synthesis. This inherent properties of neural fields allow reconstruction of “unseen” images from a learned 3D representation. Third, only one dataset (the patient’s MRI study) is needed for training,without the requirement of extensive training times and costly dataset collection(10). And fourth, the volumetric information is stored in the weights of the network, allowing improved file compression(11).

In this study a neural field super resolution approach is implemented for the task of brain MRI image enhancement. The advantages and disadvantages of the machine learning method are explored. The goal is to evaluate the potential use of neural fields as an enhancement of standard brain MRI T1 and T2 weighted series to obtain higher resolution images and improved visualizations of the neuroanatomy.

## Background

### Machine learning MRI super resolution

Maximizing MRI spatio-temporal resolution during image acquisition comes with the caveat of prolongation of study times, leading to patient discomfort and higher costs. Imaging departments with high patient volume face the need to maximize scheduling, so MRI scans are acquired in most cases in low resolution to optimize image reconstruction. Machine learning MRI super resolution methods including convolutional neural networks, latent diffusion models and generative adversarial networks have shown improvement of low resolution MRI images. All of these architectures, however, require learning a prior from a large dataset of patient studies, and they all have specific drawbacks limiting their wide adoption outside of a manufacturer closed source implementation. We refer the reader to a comprehensive reviews of the state of the art on brain MRI super resolution using deep learning methods (12).

### Neural fields

Neural networks trained to map spatial coordinates to visual information, such as images or 3D shapes, learn a parametric function of the input, storing implicitly the information in their learned parameters or weights. Multilayer perceptrons (MLP) are the most common network type used in neural fields. Since MLP have a bias toward low frequency learning, a positional encoding is implemented to enable learning of the high dimensional image features (13). Siren, as an alternative approach, implements networks with sine activation functions to improve high frequency feature learning, replacing the traditional rectified linear activation (10). Neural fields have been successfully implemented in many fields like geographic mapping, satellite imaging, robotics, and it is an ongoing research field that expanded significantly with Neural radiance fields (NerF) (14), a method that implement a multilayer perceptron and differentiable volume rendering to learn a 3D scene representation from spatial coordinates and view direction predicting color and density. NerF achieved high quality 3D novel view synthesis in a compact neural representation, training times however were high requiring more than 24 hrs of training for complex scenes. Posterior methods, such as Instant neural graphics primitives (15) achieved fast neural radiance field training using a multi resolution hash grid encoding and spherical harmonics.

## Methods

All experiments were conducted using a dedicated computer workstation with an RTX 3090 GPU (nVidia corp, Santa Monica, California) with 24 GB of VRAM. A dataset of 10 anonymized brain MRI studies including T1 and T2 weighted sequences was used for neural field training.

Relevant information about the MRI study protocol was retrieved from the Dicom metadata, including pixel image resolution, window center and width, slice thickness and voxel size.

Radscaler © (nmatik labs, Curicó, Chile)is a specialized MRI neural field training software that implements a multilayer perceptron with a high frequency positional encoding pre processing layer, allowing spatial coordinate mapping to a high dimensional feature representation to enable accurate network prediction of RGB color values for each coordinate. A supervised learning strategy was implemented for training,meaning that the neural network predicted pixel colors for each spatial coordinate and the result was optimized according to a composite loss function with regards to the ground truth image. The optimization of the network involves back propagation of gradients via the stochastic gradient descent method.

Training was conducted for 5000 epochs or 50 db peak signal to noise ratio for 5 consecutive iterations, a measure of good image quality reconstruction. Image quality was further assessed using the structure similarity index (SSIM), multi scale similarity index (MS-SSIM), and perceptual similarity index (LPIPS)(16).

Super resolution was performed using scaling factors (one up to four times original resolution) over width, height and depth dimensions. In order to maintain spatial structure, voxel size was downsized according to the new volume dimensions.

The basilar apex and the inter-thalamic adhesion were selected as region of interest (ROI) for the evaluation of image quality using peak signal to noise ratio (PSNR) with increasing zoom factors.

To assess neural interpolation capabilities for generating novel views absent from the original acquisition, intermediate images were visualized between corresponding pairs of z-voxel coordinates from the input study.

## Results

The neural field models contained 144,472,800 parameters across all processed studies, independent of input image count or dimensions. Average training time was 122.9 ± 73.9 seconds, achieving a mean PSNR of 51.3 ± 1.0 dB upon completion. Figure 1 demonstrates training progression for a T2-weighted brain MRI series over 5,000 iterations, requiring approximately 300 seconds.

**Figure 1:**
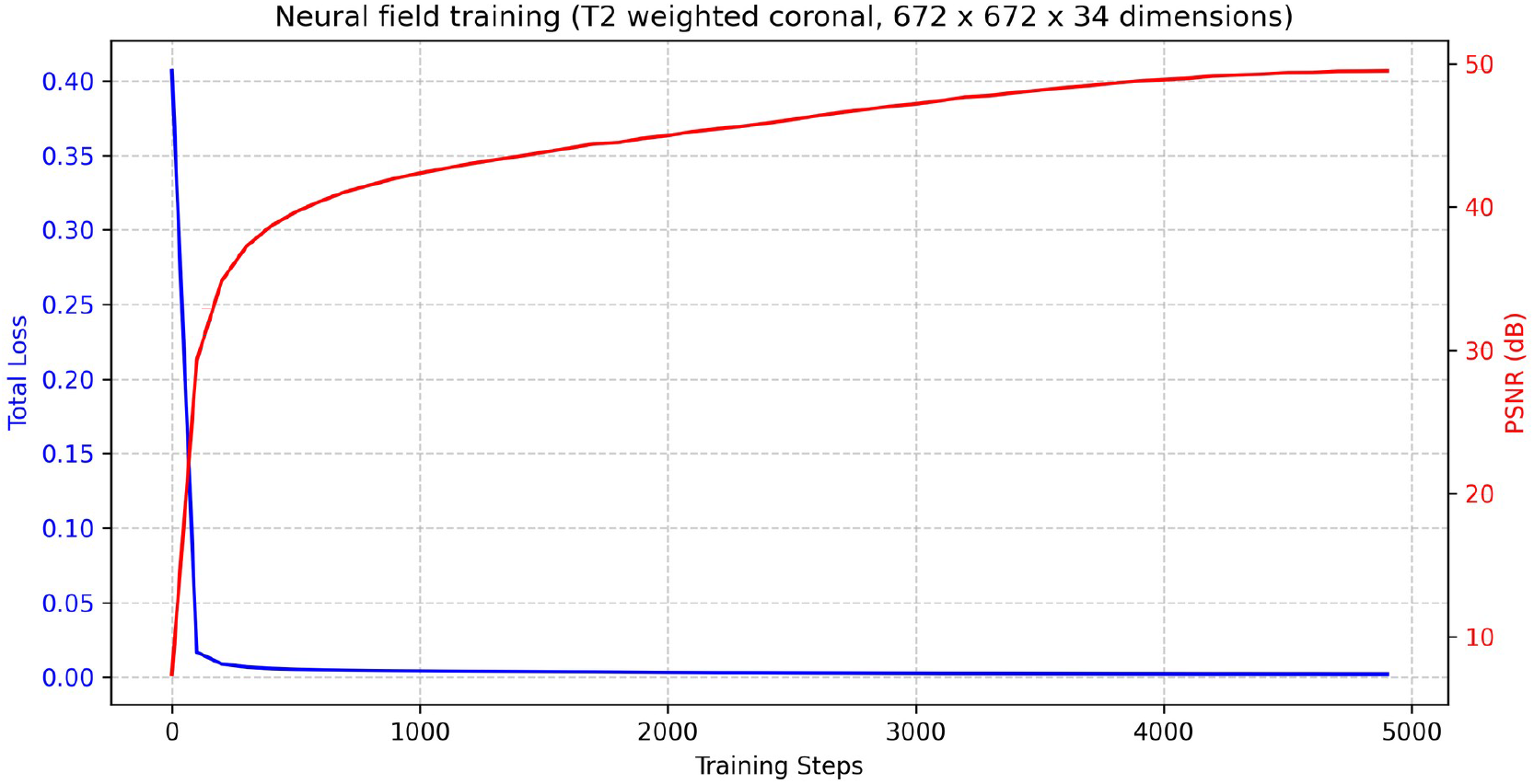
Neural field training plot.

The 10 input MRI sequences had mean image dimensions of 398.2 ± 156.8 × 439.0 ± 160.5 × 42.5 ± 34.6 pixels with corresponding voxel sizes of 0.6 ± 0.2 × 0.6 ± 0.2 × 4.5 ± 1.4 mm.

Neural field reconstruction at 2× scaling produced mean dimensions of 796.4 ± 313.6 × 878.0 ± 321.0 × 85.0 ± 69.2 pixels with voxel sizes of 0.3 ± 0.1 × 0.3 ± 0.1 × 2.2 ± 0.7 mm, representing a 50% linear dimension reduction.

At 3× scaling, reconstructions achieved 1,194.6 ± 470.4 × 1,317.0 ± 481.5 × 127.5 ± 103.8 pixels with voxel sizes of 0.2 ± 0.07 × 0.2 ± 0.07 × 1.5 ± 0.4 mm, corresponding to 66.7% linear dimension reduction.

The highest resolution 4× scaling yielded 1,628.8 ± 597.2 × 1,792.0 ± 603.1 × 160.8 ± 139.1 pixels with voxel sizes of 0.1 ± 0.05 × 0.1 ± 0.05 × 1.1 ± 0.3 mm, achieving 75% linear dimension reduction across all spatial dimensions.

Global reconstruction quality across all scaling factors demonstrated excellent performance: SSIM 0.85 ± 0.04, MS-SSIM 0.95 ± 0.01, and LPIPS 0.09 ± 0.04, indicating high-fidelity neural network predictions.

Figures 2-4 present T2-weighted coronal images centered on the basilar apex. SSIM and PSNR values in figure titles confirm robust reconstruction quality, with expected decreases at higher magnifications due to finite pixel resolution of the original 672 × 672 input (original images contain 25 × 25 pixels at maximum zoom versus 133 × 133 pixels for 5× super-resolution). Figures 5-7 demonstrate T1-weighted sagittal super-resolution with regions of interest centered on the interthalamic adhesion.

**Figure 2:**
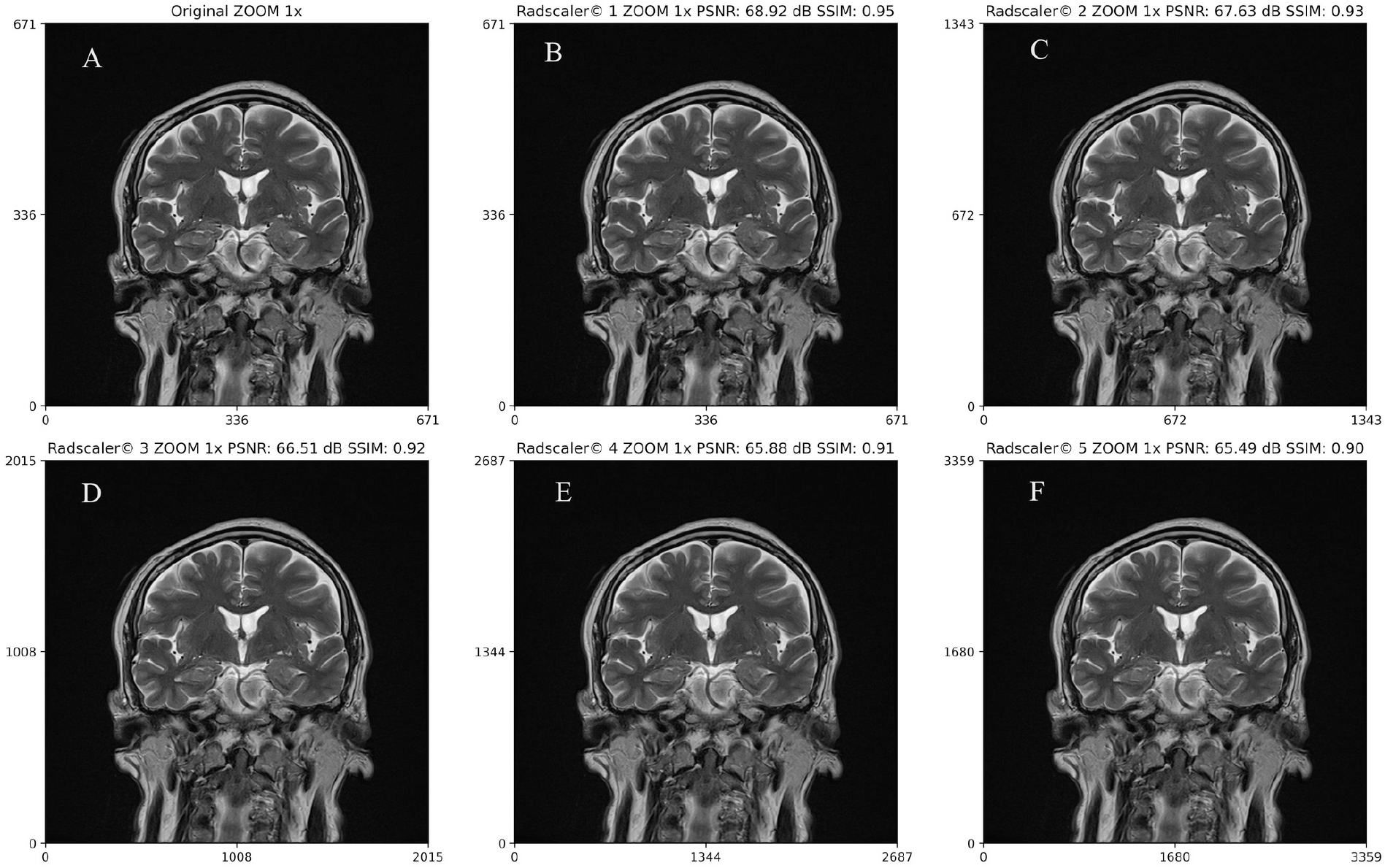

**Figure 3:**
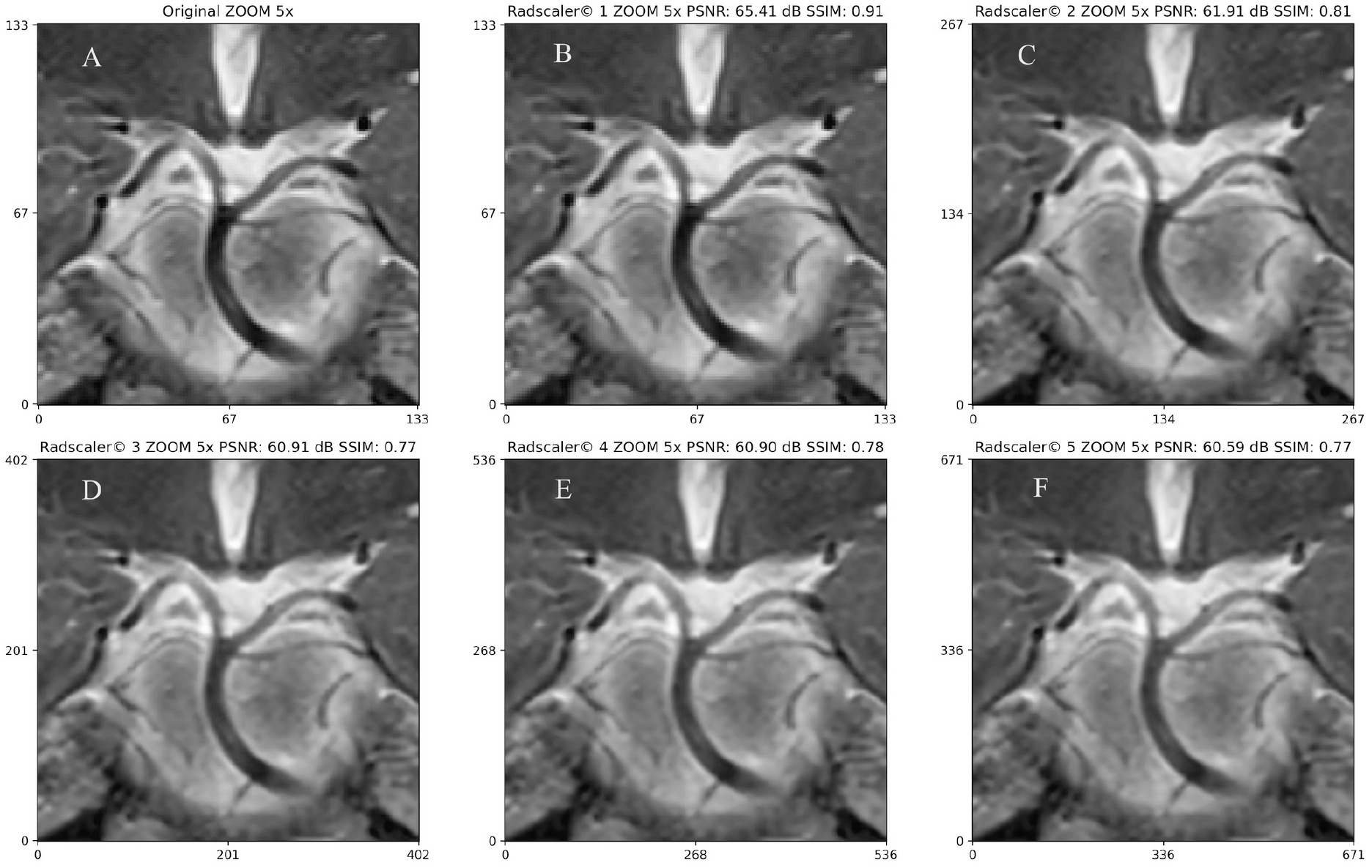

**Figure 4:**
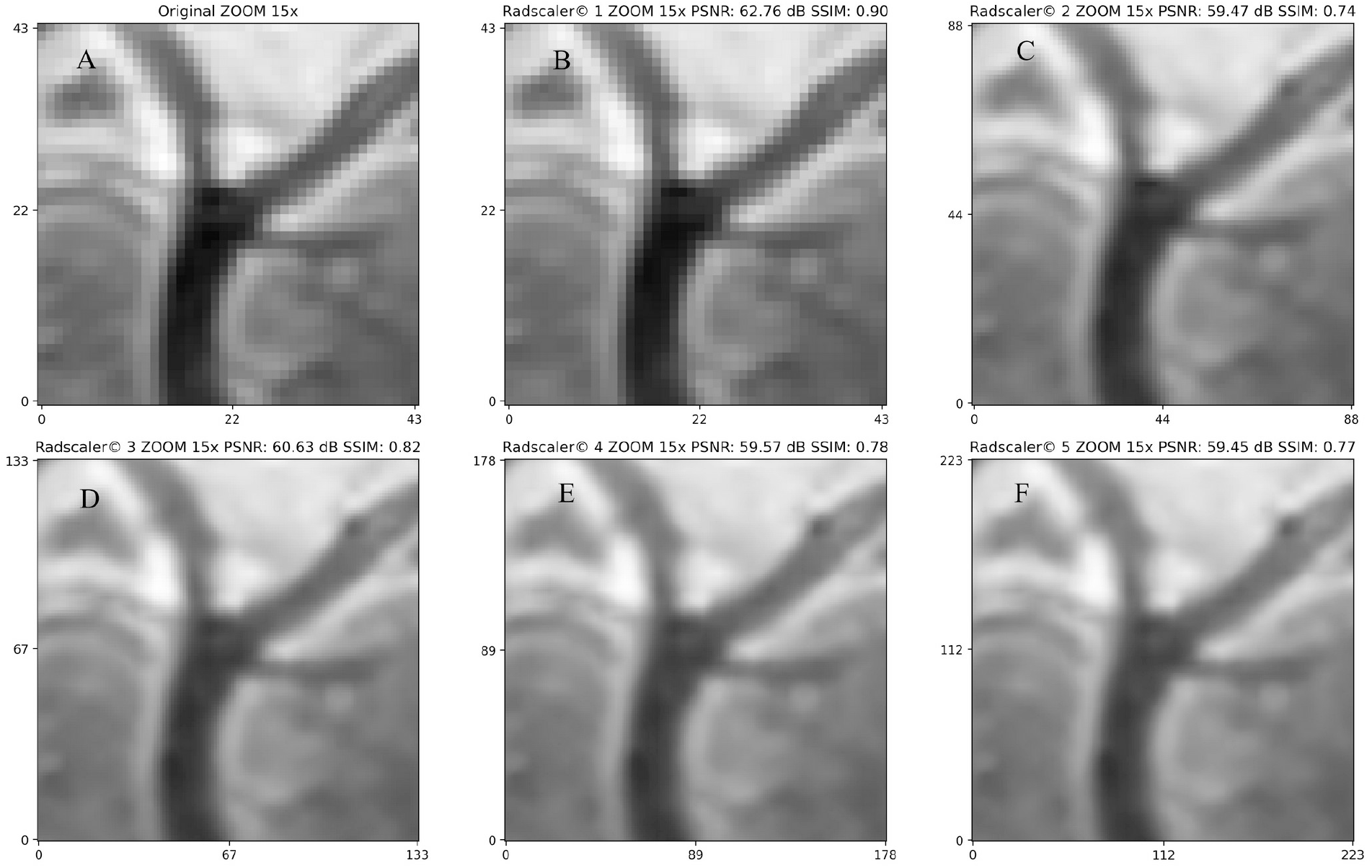

**Figure 5:**
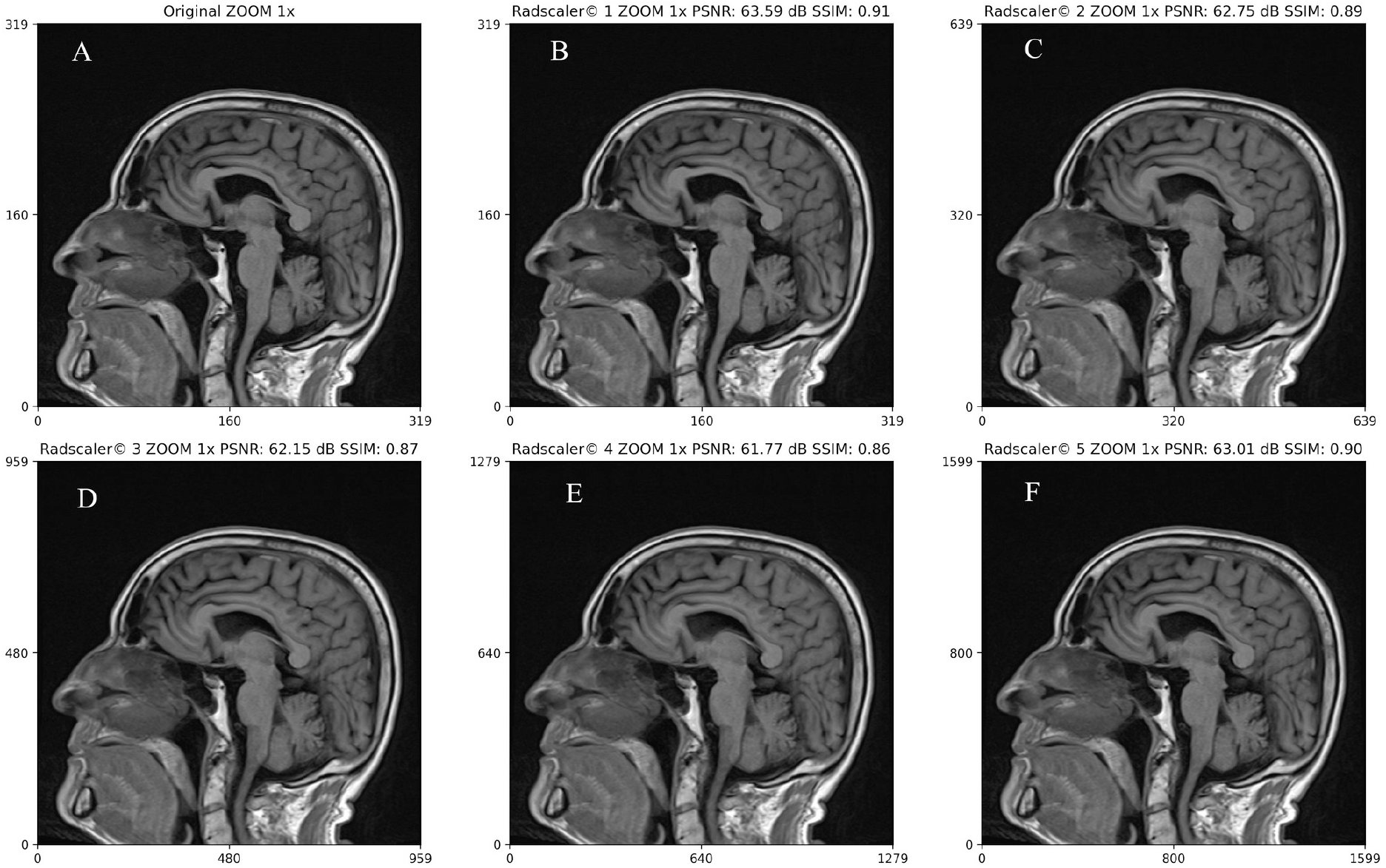

**Figure 6:**
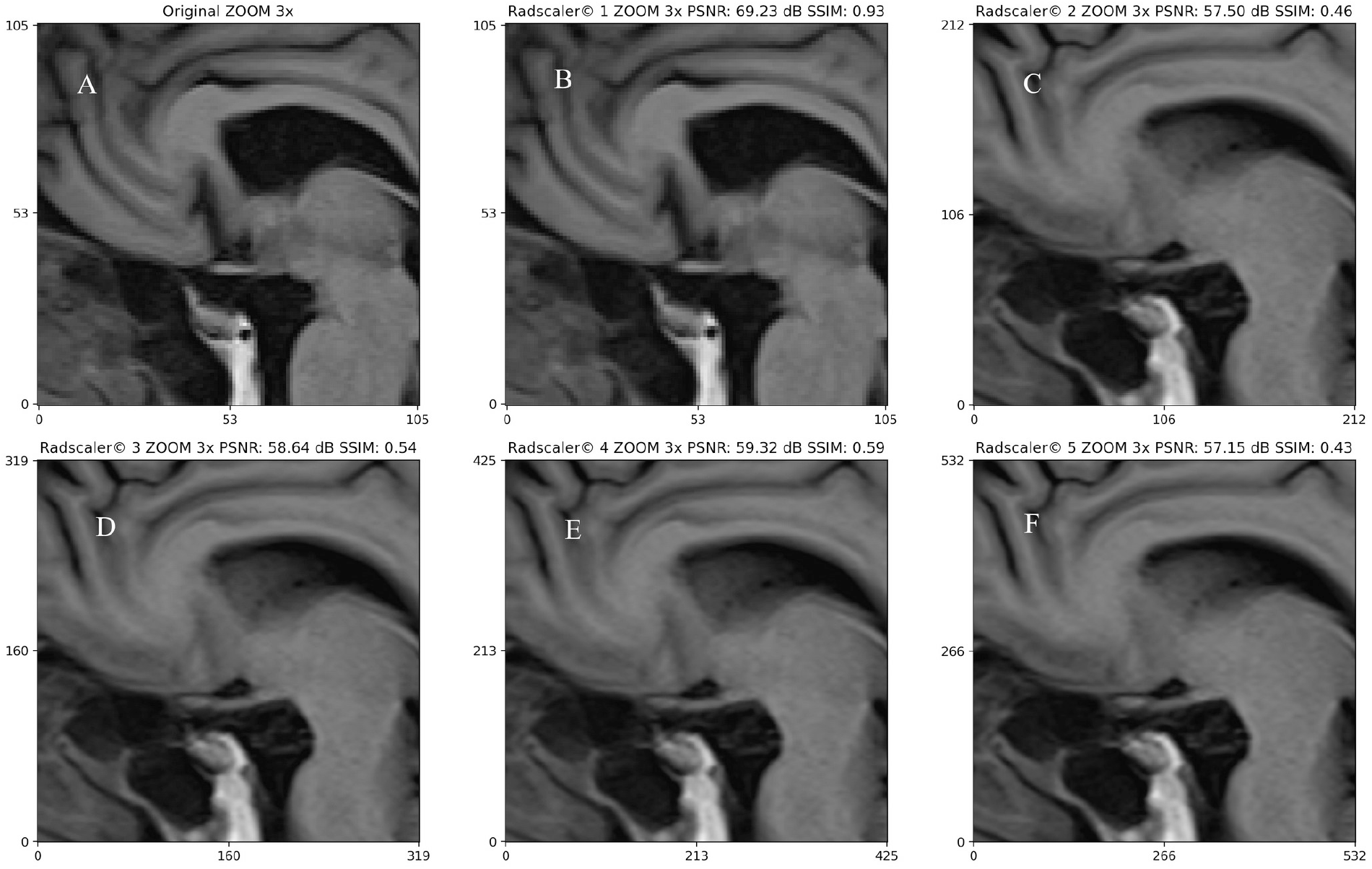

**Figure 7:**
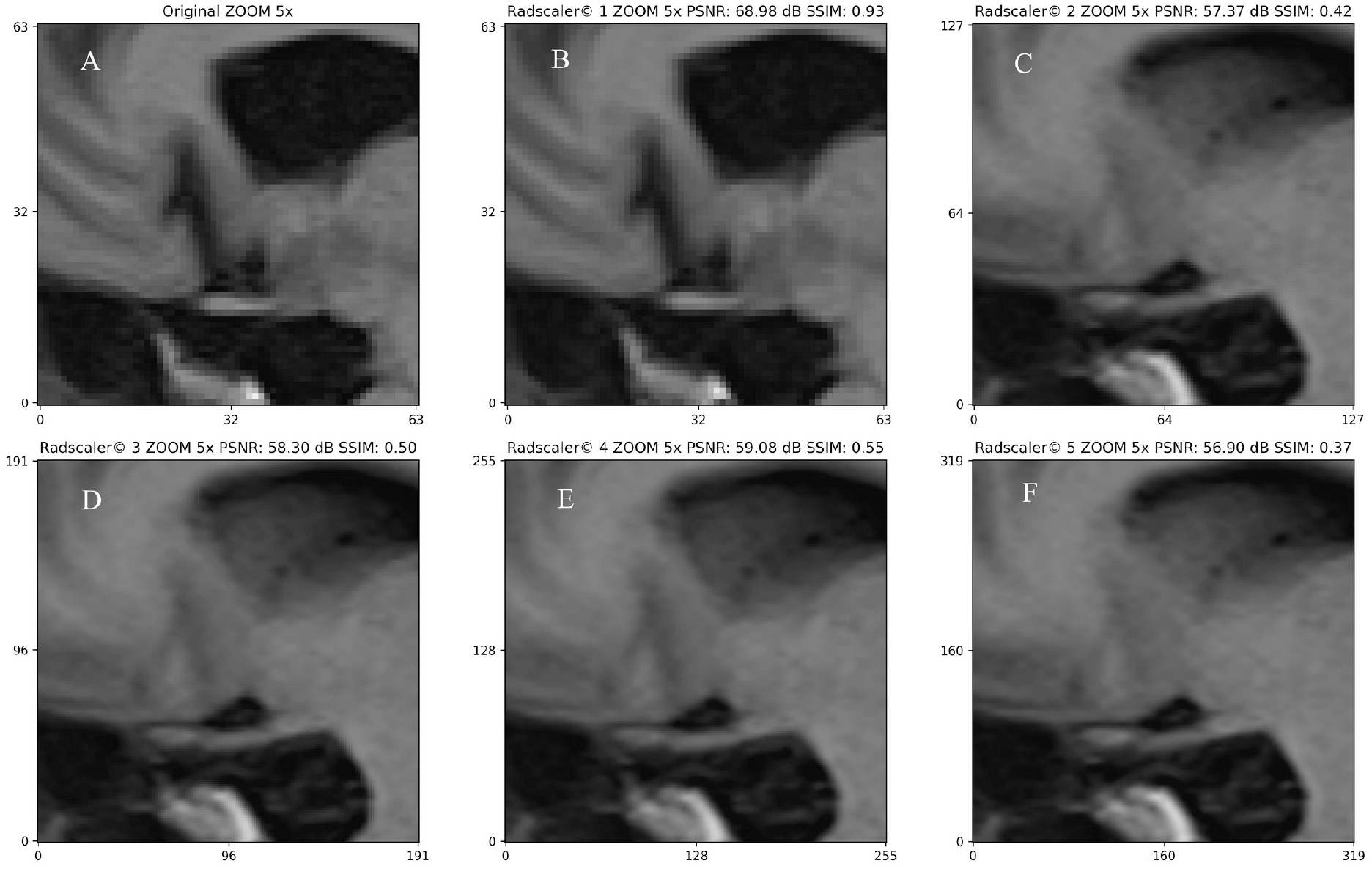

Figure 8 displays T2-weighted slices spanning z-voxel indices 9-14 from the original acquisition, while Figure 9 shows equivalent anatomical locations in the 2× SR reconstruction with world coordinate mapping. Nine intermediate slices between boundary images demonstrate neural field interpolation capabilities, synthesizing anatomically consistent views absent from the input study.

**Figure 8:**
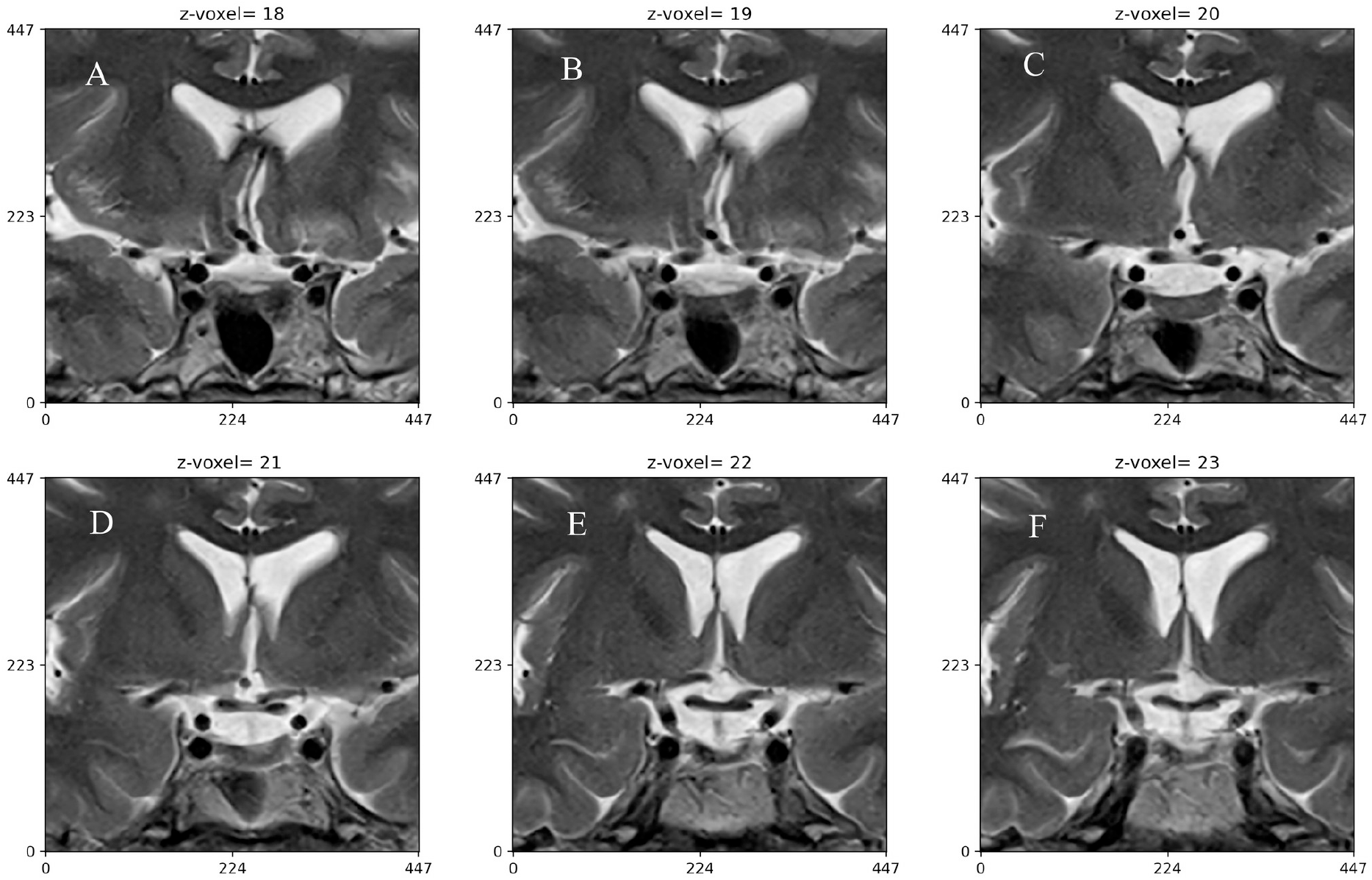

**Figure 9:**
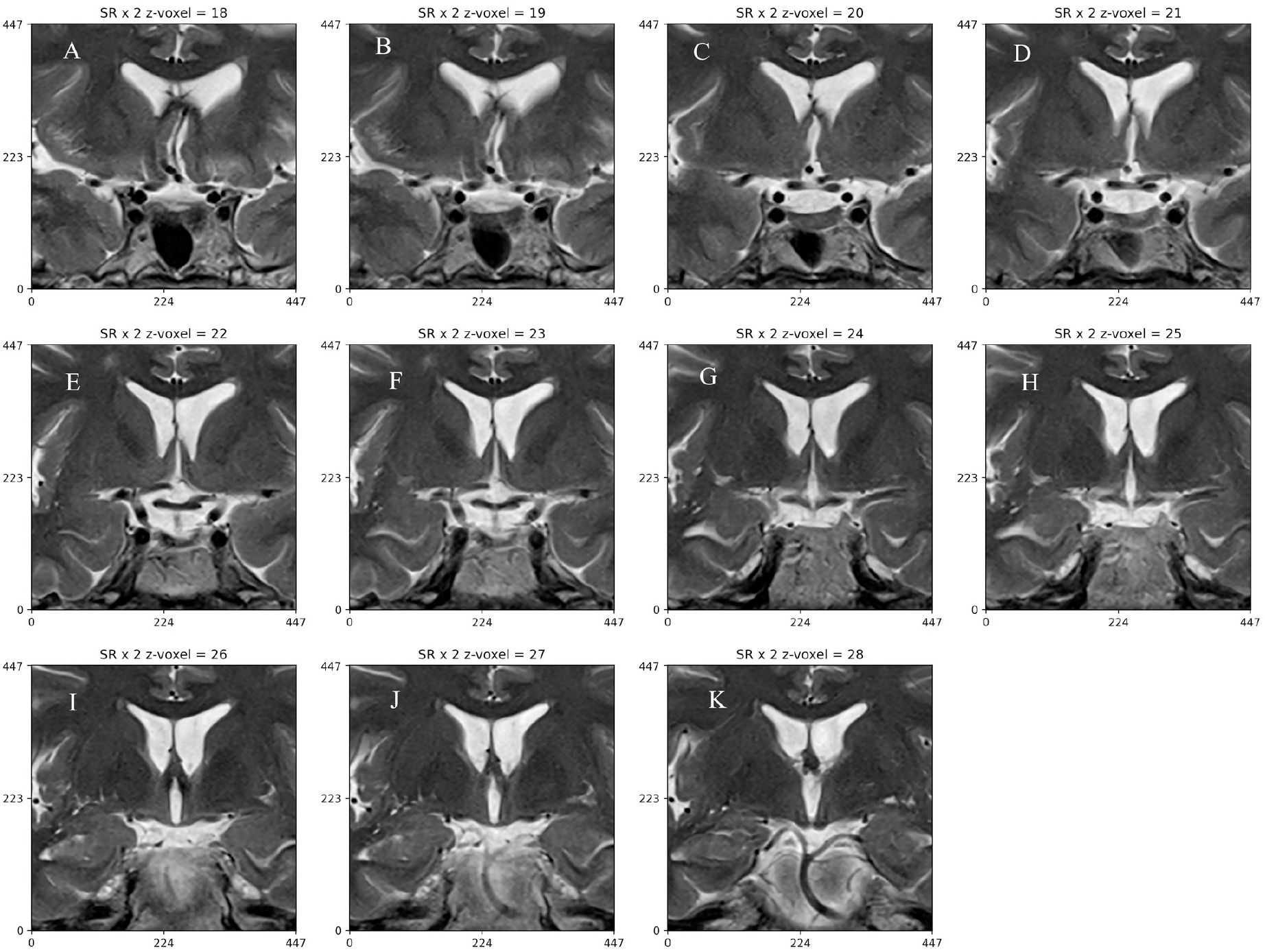

## Discussion

Brain MRI scans produce 2D slice images that collectively represent a 3D volume, typically acquired in axial, coronal, and sagittal orientations. These images are reconstructed from raw data sampled in the spatial frequency domain (k-space) using sophisticated signal processing algorithms. Image resolution is primarily determined by magnetic field strength—high-field scanners (≥3 Tesla) can achieve sub-millimeter isotropic voxel dimensions, enabling detailed anatomical visualization(17).

Beyond field strength, several acquisition parameters significantly influence image quality, including k-space matrix size, phase encoding steps, and echo times. To reduce scan duration, accelerated imaging techniques such as parallel imaging(18) or compressed sensing (19)may be employed, though these approaches can introduce spatial sampling irregularities or anisotropic voxel dimensions. Contemporary research focuses on developing advanced reconstruction algorithms, including machine learning-based approaches, to compensate for undersampled data and restore image fidelity while maintaining clinically acceptable scan times.

MRI acquisitions can exhibit either isotropic or anisotropic voxel dimensions depending on the scanning protocol. Isotropic imaging, such as a 1.5 Tesla T1-weighted sequence with 1×1×1 mm voxels, maintains uniform spatial resolution across all three dimensions, enabling consistent anatomical detail in axial, coronal, and sagittal reformations. In contrast, anisotropic sequences feature unequal voxel dimensions, where the slice thickness exceeds the in-plane resolution. This dimensional imbalance compromises spatial resolution and can degrade visualization quality when viewing or reconstructing images in planes perpendicular to the acquisition orientation.

To address these limitations, we implemented patient-specific neural field networks trained directly on individual MRI datasets, eliminating the need for large-scale training cohorts while achieving rapid convergence. State-of-the-art neural field architectures now enable high-fidelity 3D reconstruction in under five minutes (15) through several key innovations: positional encoding schemes that capture high-frequency spatial details, compact multi-layer perceptron (MLP) architectures optimized via gradient descent, and adaptive encoding functions that enhance the network’s ability to represent fine anatomical structures. This approach effectively upsamples anisotropic data to create smooth, isotropic representations suitable for multi-planar analysis and advanced visualization techniques.

Neural field-based MRI super-resolution generates novel anatomical views that extend beyond the original acquisition planes, effectively compensating for anisotropic information loss through learned 3D spatial representations. These reconstructed images represent sophisticated interpolations derived from the neural network’s continuous volumetric encoding of the patient’s original scan data, rather than synthetic content generation.

Anatomical structures, such as arteries and cranial nerves can be observe in an increased number of frames, improving spatial recognition. The obtained novel views should not be considered hallucinations or a type of generative AI, but instead a type of neural interpolation, similar in concept to the traditional re-slicing performed when implementing volume rendering of anisotropic medical images(20). The images are indeed a valuable approximation to the real patient anatomy based on real data from the patient study.

The continuous representation capabilities demonstrated by neural fields hold significant potential for advancing downstream neurosurgical applications. These include immersive virtual reality visualization for surgical planning, real-time augmented reality navigation during procedures, and high-resolution 3D printing of patient-specific anatomical models. By providing smooth, isotropic representations from anisotropic source data, neural field reconstruction bridges the gap between acquisition limitations and clinical visualization requirements.

## Conclusion

Neural fields allow high quality super resolution and novel view synthesis of brain magnetic resonance imaging. Neural field imaging processing has the potential to become a complementary source of information for neurosurgeon in the preoperative assessment of complex neurosurgical pathology.

---

**Table 1:** MRI study data.

|  | Protocol name | Repetition time (ms) | Echo Time (ms) | Phase encoding steps | Window center (intensity range) | Window width (intensity range) |
| --- | --- | --- | --- | --- | --- | --- |
| 1 | T2 TSE coronal 3mm | 4260 | 120 | 269 | 326.0 -382.3 | 712.0 – 828.3 |
| 2 | T2 FLAIR transverse 4mm | 968 | 25 | 187 | 63- 286.8 | 170.0 – 618.7 |
| 3 | T1 TSE FLAIR sagittal | 2000 | 7.4 | 230 | 401.1-526.7 | 864.0- 1104.8 |
| 4 | T2 TSE transverse 4mm | 3760 | 120 | 240 | 20.0 – 399.0 | 50.0 – 842.8 |
| 5 | T1 MPRAGE sagittal isotropic | 1930 | 2.35 | 222 | 215.2 – 342.0 | 500.4 – 734.9 |
| 6 | T2 TSE FLAIR transverse | 9000 | 96 | 168 | 169.0 – 303.9 | 394.0 – 671.9 |
| 7 | T2 TSE sagittal | 4810 | 87 | 264 | 540.0 -766.0 | 1133.0 – 1600.4 |
| 8 | T2 FLAIR transverse GRE | 694 | 21 | 211 | 165.0 -537.4 | 409.0 – 1072.0 |
| 9 | T1 MPRAGE sagittal post-Gd | 1440 | 2.78 | 270 | 457.0 – 483.9 | 948.0 – 1132.7 |
| 10 | T2 TSE coronal | 3230 | 90 | 191 | 550.0 -873.3 | 1160.0– 1773.1 |

**Table 2:** Training results.

|  | Protocol name | Training time<br>(seconds) | Training steps | PSNR<br>(dB) | Total loss |
| --- | --- | --- | --- | --- | --- |
| 1 | T2 TSE coronal<br>3mm | 299.1 | 5000 | 49.4 | 1.00e-03 |
| 2 | T2 FLAIR<br>transverse 4mm | 42.3 | 700 | 51.0 | 1.00e-03 |
| 3 | T1 TSE FLAIR<br>sagittal | 123.1 | 2000 | 51.1 | 1.00e-03 |
| 4 | T2 TSE<br>transverse 4mm | 174.7 | 2900 | 50.2 | 1.00e-03 |
| 5 | T1 MPRAGE<br>sagittal isotropic | 108.3 | 1800 | 51.7 | 1.00e-03 |
| 6 | T2 TSE FLAIR<br>transverse | 108.5 | 1800 | 52.0 | 2.00e-03 |
| 7 | T2 TSE sagittal | 144.1 | 2400 | 51.0 | 1.00e-03 |
| 8 | T2 FLAIR<br>transverse GRE | 66.5 | 1100 | 52.0 | 1.00e-03 |
| 9 | T1 MPRAGE<br>sagittal post-Gd | 54.4 | 900 | 53.4 | 1.00e-03 |
| 10 | T2 TSE coronal | 108.0 | 1800 | 51.8 | 1.00e-03 |

**Table 3:** Super resolution.

|  | Protocol name | Resolution | Dimensions ( W x H x D) | Voxel size (x, y ,z )<br>mm |
| --- | --- | --- | --- | --- |
| 1 | T2 TSE coronal 3mm | Input | $672 \times 672 \times 34$ | $0.3 \times 0.3 \times 3.9$ |
| | | SR x 2 | $1344 \times 1344 \times 68$ | $0.15 \times 0.15 \times 1.95$ |
| | | SR x 3 | $2016 \times 2016 \times 102$ | $0.10 \times 0.10 \times 1.30$ |
| | | SR x 4 | $2688 \times 2688 \times 136$ | $0.075 \times 0.075 \times 0.975$ |
| 2 | T2 FLAIR transverse<br>4mm | Input | $416 \times 512 \times 30$ | $0.4 \times 0.4 \times 5.2$ |
| | | SR x 2 | $832 \times 1024 \times 60$ | $0.2 \times 0.2 \times 2.6$ |
| | | SR x 3 | $1248 \times 1536 \times 90$ | $0.1 \times 0.1 \times 1.7$ |
| | | SR x 4 | $1664 \times 2048 \times 120$ | $0.10 \times 0.10 \times 1.30$ |
| 3 | T1 TSE FLAIR sagittal | Input | $576 \times 576 \times 30$ | $0.4 \times 0.4 \times 4.7$ |
| | | SR x 2 | $1152 \times 1152 \times 60$ | $0.2 \times 0.2 \times 2.35$ |
| | | SR x 3 | $1728 \times 1728 \times 90$ | $0.133 \times 0.133 \times 1.57$ |
| | | SR x 4 | $2304 \times 2304 \times 120$ | $0.10 \times 0.10 \times 1.18$ |
| 4 | T2 TSE transverse 4mm | Input | $600 \times 672 \times 30$ | $0.3 \times 0.3 \times 5.1$ |
| | | SR x2 | $1200 \times 1344 \times 60$ | $0.15 \times 0.15 \times 2.55$ |
| | | SR x 3 | $1800 \times 2016 \times 90$ | $0.10 \times 0.10 \times 1.70$ |
| | | SR x 4 | $2400 \times 2688 \times 120$ | $0.075 \times 0.075 \times 1.28$ |
| 5 | T1 MPRAGE sagittal<br>isotropic | Input | $320 \times 320 \times 27$ | $0.7 \times 0.7 \times 5.2$ |
| | | SR x2 | $640 \times 640 \times 54$ | $0.35 \times 0.35 \times 2.6$ |
| | | SR x 3 | $960 \times 960 \times 81$ | $0.23 \times 0.23 \times 1.73$ |
| | | SR x 4 | $1280 \times 1280 \times 108$ | $0.175 \times 0.175 \times 1.30$ |
| 6 | T2 TSE FLAIR<br>transverse | Input | $240 \times 320 \times 30$ | $0.7 \times 0.7 \times 5.2$ |
| | | SR x2 | $480 \times 640 \times 60$ | $0.35 \times 0.35 \times 2.6$ |
| | | SR x 3 | $720 \times 960 \times 90$ | $0.23 \times 0.23 \times 1.73$ |
| | | SR x 4 | $960 \times 1280 \times 120$ | $0.175 \times 0.175 \times 1.30$ |
| 7 | T2 TSE sagittal | Input | $296 \times 320 \times 21$ | $0.7 \times 0.7 \times 6.2$ |
| | | SR x2 | $592 \times 640 \times 42$ | $0.35 \times 0.35 \times 3.1$ |
| | | SR x 3 | $888 \times 960 \times 63$ | $0.23 \times 0.23 \times 2.07$ |
| | | SR x 4 | $1184 \times 1280 \times 84$ | $0.175 \times 0.175 \times 1.55$ |
| 8 | T2 FLAIR transverse<br>GRE | Input | $416 \times 512 \times 24$ | $0.4 \times 0.4 \times 5.8$ |
| | | SR x2 | $832 \times 1024 \times 48$ | $0.2 \times 0.2 \times 2.9$ |
| | | SR x 3 | $1248 \times 1536 \times 72$ | $0.133 \times 0.133 \times 1.93$ |
| | | SR x 4 | $1664 \times 2048 \times 96$ | $0.10 \times 0.10 \times 1.45$ |
| 9 | T1 MPRAGE sagittal<br>post-Gd | Input | $256 \times 256 \times 144$ | $1.02 \times 1.02 \times 1.02$ |
| | | SR x2 | $512 \times 512 \times 288$ | $0.51 \times 0.51 \times 0.51$ |
| | | SR x 3 | $768 \times 768 \times 432$ | $0.34 \times 0.34 \times 0.34$ |
| | | SR x 4 | $1024 \times 1024 \times 576$ | $0.255 \times 0.255 \times 0.255$ |
| 10 | T2 TSE coronal | Input | $280 \times 320 \times 32$ | $0.6 \times 0.6 \times 5.0$ |
| | | SR x2 | $560 \times 640 \times 64$ | $0.3 \times 0.3 \times 2.5$ |
| | | SR x 3 | $840 \times 960 \times 96$ | $0.2 \times 0.2 \times 1.67$ |
| | | SR x 4 | $1120 \times 1280 \times 128$ | $0.15 \times 0.15 \times 1.25$ |

**Table 4:** Image quality metrics.

|  | Study<br>protoc<br>ol | Scaling factor | SSIM | MS-SSIM | LPIPS<br>mean | LPIPS<br>std |
| --- | --- | --- | --- | --- | --- | --- |
| 1 | T2<br>TSE<br>coronal<br>3mm | SR x 1 | 0.85 | 0.95 | 0.08 | 0.04 |
|  |  | SR x 2 | 0.87 | 0.96 | 0.08 | 0.02 |
|  |  | SR x 3 | 0.87 | 0.96 | 0.08 | 0.02 |
|  |  | SR x 4 | 0.84 | 0.94 | 0.1 | 0.03 |
| 2 | T2<br>FLAIR<br>transve<br>rse<br>4mm | SR x 1 | 0.86 | 0.95 | 0.09 | 0.06 |
|  |  | SR x 2 | 0.85 | 0.95 | 0.1 | 0.04 |
|  |  | SR x 3 | 0.91 | 0.96 | 0.08 | 0.02 |
|  |  | SR x 4 | 0.83 | 0.94 | 0.12 | 0.04 |
| 3 | T1<br>TSE<br>FLAIR<br>sagittal | SR x 1 | 0.83 | 0.94 | 0.10 | 0.07 |
|  |  | SR x 2 | 0.83 | 0.94 | 0.1 | 0.05 |
|  |  | SR x 3 | 0.90 | 0.96 | 0.08 | 0.03 |
|  |  | SR x 4 | 0.82 | 0.93 | 0.1 | 0.03 |
| 4 | T2<br>TSE<br>transverse<br>4mm | SR x 1 | 0.86 | 0.96 | 0.08 | 0.05 |
|  |  | SR x2 | 0.87 | 0.96 | 0.08 | 0.03 |
|  |  | SR x 3 | 0.92 | 0.97 | 0.06 | 0.02 |
|  |  | SR x 4 | 0.85 | 0.95 | 0.09 | 0.03 |
| 5 | T1<br>MPRA<br>GE<br>sagittal<br>isotropic | SR x 1 | 0.81 | 0.94 | 0.09 | 0.06 |
|  |  | SR x2 | 0.80 | 0.94 | 0.1 | 0.04 |
|  |  | SR x 3 | 0.86 | 0.96 | 0.09 | 0.03 |
|  |  | SR x 4 | 0.80 | 0.93 | 0.1 | 0.04 |
| 6 | T2<br>TSE<br>FLAIR<br>transverse | SR x 1 | 0.80 | 0.94 | 0.08 | 0.05 |
|  |  | SR x2 | 0.81 | 0.9 | 0.09 | 0.04 |
|  |  | SR x 3 | 0.87 | 0.96 | 0.07 | 0.03 |
|  |  | SR x 4 | 0.80 | 0.93 | 0.1 | 0.03 |
| 7 | T2<br>TSE<br>sagittal | SR x 1 | 0.77 | 0.94 | 0.1 | 0.08 |
|  |  | SR x2 | 0.77 | 0.93 | 0.1 | 0.06 |
|  |  | SR x 3 | 0.79 | 0.94 | 0.1 | 0.05 |
|  |  | SR x 4 | 0.75 | 0.91 | 0.1 | 0.05 |
| 8 | T2<br>FLAIR<br>transverse<br>GRE | SR x 1 | 0.85 | 0.95 | 0.1 | 0.07 |
|  |  | SR x2 | 0.84 | 0.95 | 0.1 | 0.05 |
|  |  | SR x 3 | 0.88 | 0.96 | 0.1 | 0.04 |
|  |  | SR x 4 | 0.83 | 0.94 | 0.1 | 0.04 |
| 9 | T1<br>MPRA<br>GE<br>sagittal<br>post-Gd | SR x 1 | 0.89 | 0.97 | 0.03 | 0.01 |
|  |  | SR x2 | 0.90 | 0.97 | 0.04 | 0.01 |
|  |  | SR x 3 | 0.91 | 0.97 | 0.04 | 0.01 |
|  |  | SR x 4 | 0.90 | 0.97 | 0.04 | 0.01 |
| 10 | T2<br>TSE<br>coronal | SR x 1 | 0.79 | 0.93 | 0.1 | 0.06 |
|  |  | SR x 2 | 0.79 | 0.93 | 0.1 | 0.05 |
|  |  | SR x 3 | 0.87 | 0.96 | 0.08 | 0.02 |
|  |  | SR x 4 | 0.77 | 0.92 | 0.1 | 0.05 |

